# Neural correlates of response inhibition and performance monitoring in binge watching

**DOI:** 10.1101/868984

**Authors:** Carolin Kilian, Kyra Luisa Bröckel, Rebecca Overmeyer, Raoul Dieterich, Tanja Endrass

## Abstract

With the increasing popularity of internet streaming portals, the question why people develop binge-watching behavior has become a focus of scientific research and its addictive potential is discussed. The current study examined neural correlates of binge-watching during inhibition in a go/nogo task and performance monitoring using electroencephalography. Participants reported high binge-watching behavior (HBW, n = 35) or no binge-watching (NBW, n = 33) episode during the past four weeks. Compared to the NBW group, HBW showed larger P3 during response inhibition and larger error-related negativity (ERN) for errors in the flanker task. Group differences in behavioral measures were not observed. The current results suggest that binge watching may be related to both (1) increased neural recruitment during response inhibition as indicated by the increased P3 to facilitate normal inhibitory performance and (2) enhanced performance monitoring as indicated by the increased ERN. As this neurocognitive profile differs from observations in other addictive and excessive behaviors, implications for this budding field are discussed.

## Introduction

With the increasing popularity of internet streaming portals, binge watching (BW) has become a widely common behavior and moreover an area of scientific interest. BW is usually defined by the number of TV shows watched in one session (e.g. Trouleau et al., 2016; Walton-Pattison et al., 2018). Based on statistical modelling, Trouleau and colleagues identified a cut-off of watching 2 episodes in one session to define a BW session regardless of the length of TV show, the day of the week, and the medium used. Up to now, only few theories have attempted to explain why people indulge in watching TV Shows (Panda & Pandey, 2017; Walton-Pattison et al., 2018). One example is the description of a vicious circle, in which BW serves as a short-term maladaptive coping strategy to avoid unpleasant feelings and daily life problems (Panda & Pandey, 2017). However, BW as avoidance behavior may increase negative states and problems which in turn leads to further BW. Furthermore, BW has been associated with automaticity, anticipated regret of engaging in binging, and goal-conflicts (Walton-Pattison et al., 2018). Automaticity has been described as an impulsive process with integrated reward mechanisms that will transform watching series from a goal-directed behavior to automatic or habitual behavior. Initial studies reported a relationship between BW and impulsivity (Orosz et al., 2016; Riddle et al., 2018), which is a prominent risk factor for excessive and addictive behaviors such as internet gaming disorder (e.g. Ko et al., 2014). Both mechanisms, vicious circle and automaticity, are well known from addiction research (Koob & Volkow, 2010; Romer Thomsen et al., 2014) and the addictive potential of BW is a matter of current debate, warranting studies on processes facilitating goal-directed behavior.

Goal-directed behavior depends on cognitive control, which is compromised in addictive disorders and other mental disorders (Goschke, 2014). Cognitive control functions, such as response inhibition and performance monitoring, play a key role in understanding mental disorders. In this paper, neural correlates of response inhibition and performance monitoring in BW are investigated and may help delineate impulsive and compulsive aspects of BW. The ability to stop or withhold inappropriate responses is an important aspect of cognitive control and facilitates self-regulation and goal-directed behavior (Bari & Robbins, 2013). Response inhibition is commonly measured using go/nogo tasks, in which participants have to respond quickly to frequently occurring go stimuli but also have to suppress responses to rare nogo stimuli. Reduced nogo accuracy has been observed for internet gaming disorder and excessive internet use (Kim et al., 2017; Li et al., 2019; Littel et al., 2012; Zhou et al., 2010). However, to account for potential speed-accuracy trade-off, it has been suggested to use the weighted response accuracy in relation to the reaction time as a behavioral correlate (Liesefeld & Janczyk, 2018).

Response inhibition has been investigated with the electroencephalogram (EEG) and inhibitory processes have been related to event-related potentials (ERP) following nogo stimuli: a frontocentral negativity (N2) after 200-300 ms, a frontocentral positivity (P3a) after 250 – 400 ms, and a parietal positivity (P3b) starting at 300 ms (Folstein & Van Petten, 2008; Polich, 2007). While the N2 is associated with conflict detection, the P3 complex is interpreted as a neural marker for actual motor inhibition: P3a is associated with attention and orientation and P3b with suppression of the former reaction (Enriquez-Geppert et al., 2010; Polich, 2007). Whereas substance use disorders seem to be quite robustly characterized by hypoactive neural correlates of response inhibition (for systematic review see Luijten et al., 2014), the evidence regarding alterations of N2 and P3 amplitudes in addictive behaviors is inconclusive: the N2 appears to be unaffected in internet gaming disorder (Kim et al., 2017; Li et al., 2019; Littel et al., 2012), but increased in excessive smartphone use (Chen et al., 2016; Gao et al., 2019), and reduced in excessive internet use (Dong et al., 2010; Zhou et al., 2010). The P3 was shown to be reduced in internet gaming disorder and excessive smartphone use (Gao et al., 2019; Li et al., 2019) but increased in excessive internet use (Dong et al., 2010). However, these P3 findings were neither replicated for internet gaming (Kim et al., 2017; Littel et al., 2012) nor excessive smartphone use (Chen et al., 2016).

Performance monitoring reflects the ability to flexibly adapt actions responding to environmental changes or performance problems (Ullsperger et al., 2014). Therefore, context, progression, and results of actions must be monitored continuously, and appropriate autonomous, cognitive, or affective adaptations must be derived. A valid and reliable neural marker for error-processing in the response-locked EEG is the error-related negativity (ERN; Gehring et al., 1993; Ullsperger et al., 2014). The ERN is characterized by a frontocentral negativity, which appears 50-100 ms after an error in speeded choice tasks and is generated by the posterior medial frontal cortex. First studies have examined the ERN in addictive behaviors such as internet gaming disorder (Littel et al., 2012), internet use (Zhou et al., 2013) and “food addiction” (Franken et al., 2018). In internet gaming disorder, which has been acknowledged as behavioral addiction by the World Health Organization since the 11^th^ revision of the International Classification of Disease, the ERN reduction was correlated with weekly hours of gaming (Littel et al., 2012). The reduction of ERN amplitudes has further been observed in substance use disorders and suggested as a potential endophenotype (Euser et al., 2013; Luijten et al., 2014). A recent meta-analysis concluded that the reduction of ERN may reliably dissociate internalizing vs. externalizing disorders and could represent a marker for this dissociation (Pasion & Barbosa, 2019). Furthermore, the ERN is influenced by motivational factors such as reward and punishment as a function of task performance or task context (Endrass et al., 2010; Hajcak et al., 2005; Potts, 2011). Larger ERNs for punished errors were observed in persons with high threat sensitivity (Potts et al., 2006), whereas the ERN was found to be enhanced in a reward context in smokers (Schlienz et al., 2013).

Despite previous findings suggesting an addictive or generally pathological potential of BW, BW has not been sufficiently characterized in terms of behavioral and neural alterations in cognitive control. However, in a relatively young field, this is essential to facilitate model development and integration with other excessive or addictive behaviors. Thus, this paper examines neural correlates of response inhibition and error processing in individuals showing excessive binge watching (high binge watcher, HBW) in contrast to individuals who do not engage in binge watching (NBW). Regarding the aforementioned findings on behavioral addiction and response inhibition, the HBW group should show performance deficits in the go/nogo task compared to the NBW group. Further, based on findings for addictive behaviors, we expected alterations of N2, P3a, and P3b amplitudes for HBW vs. NBW groups, but refrained from formulating directional hypotheses due to inconclusive prior results. With respect to performance monitoring, we assumed that the HBW group would show a reduced frontocentral ERN in contrast to NBW group. In addition, the incentive context was varied between trials, in that participants could earn points (gain condition) or avoid losing points (lose condition) if responses were executed correctly and fast. The group difference should be influenced by the incentive context, which would result in an interaction between group (NBW, HBW) and context (gain, loss avoidance).

## Materials and methods

### Participants

Thirty-five participants were included in the HBW group and the NBW group comprised 33 participants (Table 1). All Individuals were recruited via local advertisement and were screened with an online pre-survey and a short telephone interview. Exclusion criteria were known neurological disorders, previous head trauma, intellectual disability, lifetime diagnoses of schizophrenia, psychotic disorder, and/or manic episode; current diagnoses of eating disorder, depressive episode, and previous or current high alcohol and/or drug consumption.

**Table 1.** Demographic, clinical and performance data of NBW and HBW group (mean and standard deviations are reported).

|  | *n* | NBW | HBW | *F/X*<sup>2</sup> | *p* |
| --- | --- | --- | --- | --- | --- |
| **Demographic** |  |  |  |  |  |
| Gender (female/male) | 68 | 18/15 | 22/13 | 0.48 | .49 |
| Education (secondary school/ high school) | 67 | 2/31 | 4/30 | 0.67 | .41 |
| Age (years) | 68 | 25.97 (6.30) | 25.46 (5.57) | 0.14 | .71 |
| **Mean (SD) Watching behavior** |  |  |  |  |  |
| BW sessions | 67 | 0.15 (0.71) | 15.53 (7.44) | 139.54 | <.001 |
| Episodes per session | 66 | 1.91 (3.59) | 3.94 (2.52) | 7.19 | .01 |
| WLOC | 68 | 3.79 (4.74) | 13.06 (5.22) | 58.47 | <.001 |
| **Mean (SD) clinical score** |  |  |  |  |  |
| BIS-11 | 68 | 56.34 (7.91) | 60.34 (8.23) | 4.10 | .05 |
| DASS Anxiety | 68 | 1.18 (1.99) | 1.46 (1.31) | 0.46 | .50 |
| OCI-R | 60 | 11.60 (9.79) | 13.66 (7.91) | 0.81 | .37 |
| **Mean (SD) performance data** |  |  |  |  |  |
| Go/nogo |  |  |  |  |  |
| Nogo Accuracy (%) | 68 | 86.7 (12.0) | 86.7 (12.5) | <0.01 | .99 |
| RT (ms) | 68 | 286 (27) | 283 (32) | 0.03 | .85 |
| BIS | 68 | -0.05 (0.94) | 0.05 (1.10) | 0.38 | .71 |
| MIFLAT |  |  |  |  |  |
| Accuracy (%) | 64 | 84.36 (8.67) | 86.00 (6.40) | 0.75 | .39 |
| RT error (ms) | 64 | 262.70 (25.58) | 257.00 (21.34) | 0.94 | .34 |
| RT correct (ms) | 64 | 328.25 (31.30) | 327.18 (29.78) | 0.02 | .89 |
| Post-error slowing (ms) | 64 | 7.73 (15.28) | 9.51 (8.49) | 0.34 | .57 |

*Abbreviations.* NBW, No Binge Watching; HBW, High Binge Watching; BW, Binge Watching; WLOC, Watching Loss of Control; BIS-11, Barratt Impulsiveness Scale; DASS, Depression Anxiety Stress Scales; OCI-R, Obsessive-Compulsive Inventory-Revised; MIFLAT, monetary incentive flanker task. Degrees of freedom: $df = n - 2$ .

For assessment of individual watching behavior, all participants reported frequency and duration of their BW episodes in the previous four weeks. A BW episode was defined as watching more than three episodes of TV-shows in one session. Individuals in the HBW group reported at least ten BW episodes in the previous four weeks while individuals in the NBW group reported no BW episodes in the same period. The BW definition used in this study differs from those usually applied for three reasons: First, we aimed to include individuals with recurrent BW to represent a compulsive instead of situational behavior (e.g. watching series at weekends only). Second, we supposed that our sampling procedure (i.e. convenience sample) will overestimate the number of young persons aged between 20 and 30, a subgroup that reports BW behavior more frequently than older adults (Sung et al., 2018). Third, we pilot-tested the BW assessment in a sample of 159 adults aged 18 to 65 years. In the sample, 39.6% reported to watch more than 2 episodes in one session at least once during the past four weeks whereof 20.1% were watching more than 3 episodes in one session. Since we were interested in individuals with problematic watching behavior, we decided to use the cut-off of watching more than 3 episodes in one session. It should be mentioned that at the time the study was carried out, no validated measure assessing symptoms of problematic BW was available. Impulsivity was assessed with the Barratt-Impulsiveness-Scale (BIS-11; Patton et al., 1995). Compulsive symptoms were evaluated using the Obsessive-Compulsive Inventory-Revised (OCI-R; Foa et al., 2002) and anxiety symptoms using the Depression Anxiety Stress Scales (DASS; Henry & Crawford, 2005).

The study was conducted in accordance with the ethical guidelines of the Declaration of Helsinki and approved by the local ethics committee of the TU Dresden (EK 465112018). Participants received verbal and written information about the procedure and content of the study and gave written informed consent.

#### Tasks and Procedure

Participants completed two different tasks during electroencephalography.

#### Go/nogo task

The task consisted of 12 practice trials and two blocks of 128 trials. During the task a white circle presented on a grey background was always visible. At the center of the circle either a green square was presented as go stimulus (75 % of all trials) and participants should respond as quickly as possible with their right index finger, or a red square was shown as nogo stimulus (25 % of all trials) and participants had to withhold their response. Stimuli were presented for 500 ms and were separated by a variable inter-stimulus interval of 900 to 1200 ms (mean: 1150 ms). Nogo trials could immediately follow each other or were separated with up to 5 go trials. Completion of the task took about eight minutes.

#### Monetary incentive flanker task (MIFLAT)

A modified arrow-version of the flanker task (e.g. Endrass et al., 2010) was used to measure the ERN (Figure 1). The motivational context was varied between trials. In the gain condition, fast and correct responses were rewarded (40 points) while errors resulted in a reward omission (0 points). In the loss condition, errors and slow responses were punished (minus 40 points) and fast and correct responses resulted in punishment omission (0 points). The trial started with an initial fixation cross surrounded by a green or red frame (for 500 ms), indicating a gain or loss context (50 % of all trials), respectively. Then, four vertically arranged arrows were presented within the frame and after 100 ms the target arrow was presented in the center for 30 ms. Half of the trials were compatible, i.e. all five arrows pointed in the same direction, and the remaining trials were incompatible, i.e. the target and flanker arrows pointed in opposite directions. Incentive context, compatibility, and target direction were presented in pseudo-random order and balanced between conditions. Participants were instructed to indicate the direction of the target arrow with a left or right button press as quickly and as accurately as possible. After a response interval of 900 ms following target onset or 600 ms after the response, a performance feedback was presented for 800 ms. Depending on correctness and response time, participants could earn or lose 40 points in the gain or loss condition. If their performance was incorrect or slower than an adaptive deadline based on their response time, rewards were omitted, or 40 points were lost. The response deadline was calculated based on performance and reaction time in order to obtain a rate of 20% negative feedbacks for each context. Participants could earn a bonus of up to 5 EUR. The task included 640 trials with a duration of 2.53 to 2.75 s and lasted about 28 minutes.

**Figure 1.**
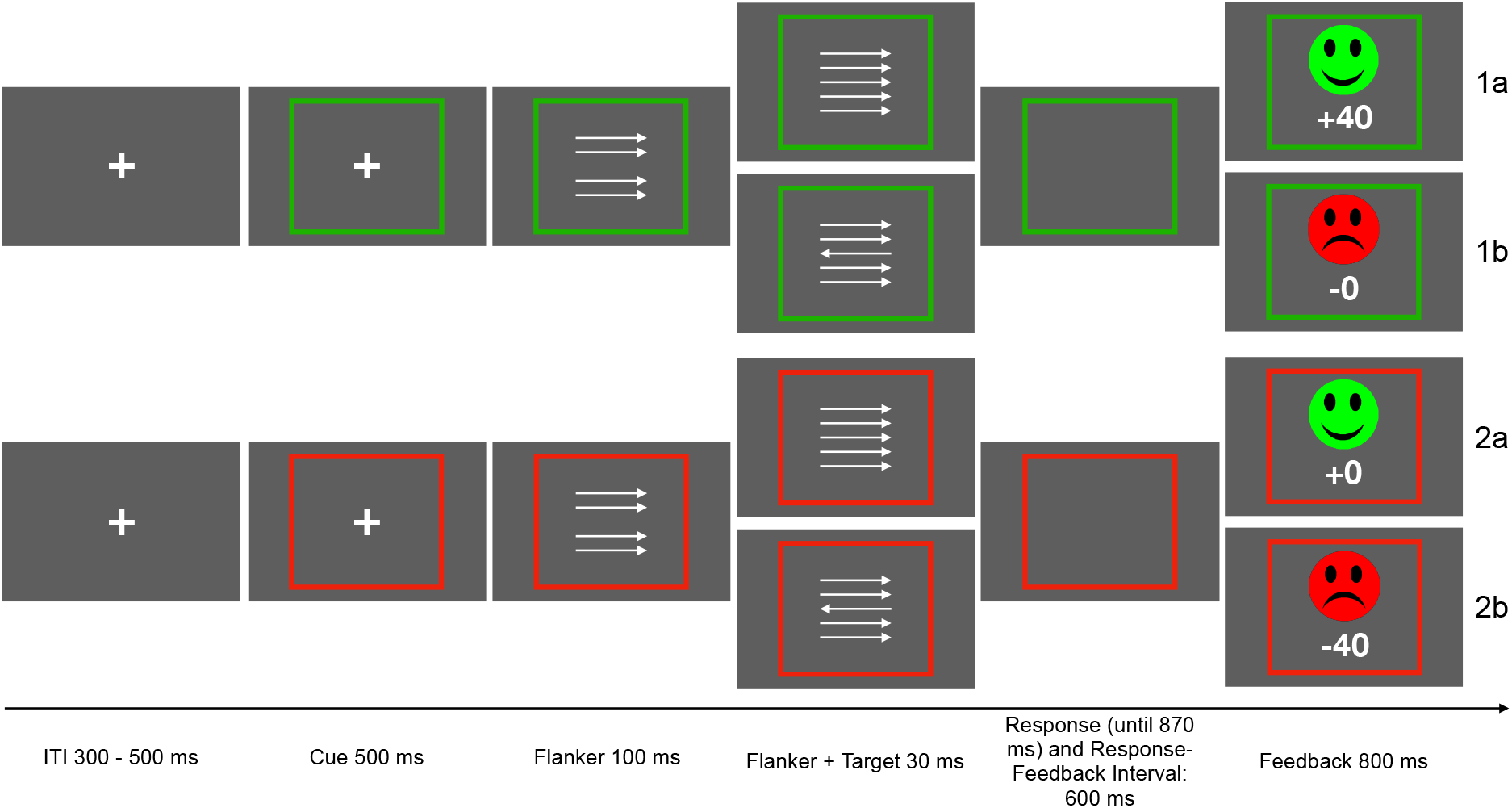
Schematic depiction of the monetary incentive flanker task (MIFLAT). The gain condition is displayed in the upper, the loss condition in the lower row.

#### Electroencephalogram recording and data analysis

The EEG was recorded with 64 Ag/AgCl electrodes using two 32-channel BrainAmp amplifiers (Brain Products GmbH, Munich, Germany) and an EasyCap electrode cap with equidistant electrode locations (EasyCap GmbH, Herrsching-Breitbrunn, Germany). Impedances were kept below 5 kOhm during the recording and the ground and reference electrode were mounted next to Fz. Below the left and right eye two electrodes were placed for capturing eye movements in combination with electrodes mounted in the cap. Data were continuously registered at 500 Hz sampling frequency.

Data were preprocessed and analyzed using EEGLAB 14.1 (Delorme & Makeig, 2004). The EEG was re-referenced to common average reference and bandpass filtered with 0.5 to 30 Hz. Ocular artifacts were removed using an infomax independent component analysis as implemented in EEGLAB. Epochs spanning from 200 ms before to 800 ms after stimulus onset in the go/nogo task and after response onset in the MIFLAT were obtained. Epochs that contained deviations greater than 5 SDs of the mean probability distribution were automatically rejected, as were epochs with early (< 100 ms in go/nogo; < 150 ms in MIFLAT) or late responses (> 600 ms). On average, *n* = 5.7 and *n* = 7.7 trials were excluded in the go/nogo and MIFLAT task, respectively. Groups did not differ with regard to the number of trials excluded due to reaction time in the go/nogo (go, *F*(1,67) = .27, *p* = .61; nogo, *F*(1,67) = .63, *p* = .43) or the MIFLAT (correct, *F*(1,63) = .12, *p* = .73; error, *F*(1,63) = 1.26, *p* = .27) task. Individual and group averages were calculated as stimulus-locked averages for correct go and nogo trials in the go/nogo task and as response-locked averages for incorrect responses in the gain and loss context of the MIFLAT. EEG activity during the time window of 200 ms before stimulus or response onset served as baseline and was subtracted from the data.

All statistical analyses were performed with SPSS (IBM Corp., 2017) and MATLAB (The MathWorks, Inc., 2017). In the go/nogo task, stimulus-locked ERPs (N2, P3a, and P3b) were analyzed for correct go and nogo trials. The N2 and P3a were determined in a frontocentral electrode cluster (FCz, FC1, FC2, Cz). The N2 was defined as the most negative value in the time window 230 to 270 ms after stimulus onset and P3a as the average amplitude from 300 to 380 ms after stimulus onset. The P3b was obtained as average amplitude in the time window from 320 to 450 ms in a parietal electrode cluster (CPz, CP1, CP2, Pz, P1, P2). Clusters were chosen based on the topography of the respective component (Huster et al., 2013; Polich, 2007) and were used in studies examining response inhibition in substance use disorder or behavioral addiction (Littel et al., 2012; Luijten et al., 2011). In MIFLAT, the mean amplitudes for correct and incorrect response trials were quantified as mean amplitude at frontocentral electrodes (FCz, Cz), where the ERN was most pronounced, in the time window centered around the grand-average ERN peak (M = 50 ± 10 ms). Amplitudes were determined for incompatible trials only as only few errors were committed on compatible trials (< 1%). For incorrect responses, on average n = 40 trials (*Min* = 8; *SD* = 20) were included for the loss context and n = 49 trials (*Min* = 9, *SD* = 23) in the gain context. Groups did not differ regarding the number of included trials, *t*(62) < 1.0, *p* > .3. The minimum number of error trials exceeded the previously published recommendation of six trials to reliably measure the ERN (Olvet & Hajcak, 2009) and only two participants had less than ten trials in any condition (Fischer et al., 2017).

Demographic and clinical data were compared between groups (HBW vs. NBW) using one-way analyses of variance (ANOVAs). In the go/nogo task, inhibitory performance was measured as the proportion of correct inhibitions in nogo trials, reaction time in go trials, and the difference between the standardized accuracy minus the mean go reaction time (Balanced-Integration-Score, BIS, Liesefeld & Janczyk, 2018) and compared between groups with one-way ANOVAs. ERP amplitudes were analyzed with repeated measures ANOVAs with the within-subject factor trial type (go, nogo) and the between-subject factor group (NBW, HBW). In the MIFLAT, accuracy (proportion of correct responses), error and correct RT were quantified depending on stimulus compatibility and reward context. Post-error slowing was defined as the difference of mean RTs in correct trials following errors minus RTs in correct trials following correct responses depending on context. Repeated measures ANOVAs were conducted for each of the following dependent variable: response accuracy, RT, and ERP amplitude. Context (gain vs. loss), compatibility (compatible vs. incompatible), correctness (correct vs. incorrect) served as the within-subject factors, and group (HBW vs. NBW) as between-subject factor. A potential association between ERN amplitudes and BW sessions within the HBW group was examined using Pearson correlations.

In the MIFLAT, four participants were excluded due to of insufficient response accuracy (< 38.6 % errors, *n* = 2), termination after two blocks (*n* = 1) and an outlier analysis of the EEG data (*n* = 1). Individuals with missing data (educational level: *n* = 1; OCI-R: *n* = 8) or implausible data (number of BW sessions = 125: *n* = 1; number of episodes watching in one BW session was equal to the duration of one BW session: *n* = 2) were only excluded for the respective analyses to ensure the power of ERP analyses.

## Results

### Demographic and behavioral data

Groups did not differ with respect to age, education, gender, DASS-or OCI-R-scores (Table 1). Compared to NBW, the HBW group reported significantly more BW sessions, *F*(1,67) = 139.5, *p* = < .001, *η*^2^_p_ = .68, more episodes during one session, *F*(1,67) = 7.2, *p* = .01, η^2^_p_ = .10, and more subjective loss of control, *F*(1,67) = 58.5, *p* = < .001, η^2^_p_ = .47. BIS-11-scores were also significantly higher in the HBW group in contrast to NBW, *F*(1,67) = 4.10, *p* = .048, η^2^_p_ = .06. Group differences in behavioral data were neither observed in the go/nogo task nor in the MIFLAT (see Table 1).

MIFLAT: Fewer errors, *F*(1,62) = 244.58, *p* < .001, *η^2^_p_* = 0.80, and faster correct RTs, *F*(1,62) = 1983.0, *p* < .001, *η^2^_p_* = 0.97, were observed for compatible than for incompatible trials. RTs were shorter for incorrect than correct responses, *F*(1,62) = 3972.5, *p* < .001, *η*^*2*^_*p*_ = .99. Performance also varied as a function of compatibly and context. Incompatible errors were less frequent and response times were longer in the loss compared to the gain condition, as indicated by the interactions of context and compatibility, *F*(1,62) = 59.8, *p* < .001, *η*^*2*^_*p*_ = .49 and *F*(1,62) = 12.9, *p* < .001, *η*^*2*^_*p*_ = .17. Significant group main effects or interactions with experimental factors were not observed.

### Neural response: response inhibition

Group averages and topographies are displayed in Figure 2. N2 was more pronounced after nogo than after go stimuli, *F*(1,66) = 118.8, *p* < .001, η^2^_p_ = .64. Neither a group difference, *F*(1,66) < 0.1, *p* = .91, *η*^2^_p_ < .01, nor an interaction effect, *F*(1,66) < 0.1, *p* = .94, *η*^2^_p_ < .01, was observed for the N2. The P3a was larger in nogo trials than in go trials, *F*(1,66) = 192.8, *p* < .001, *η*^2^_p_ = .75. For the P3a a significant main effect of group was observed: P3a was more pronounced in HBW than in NBW, *F*(1,66) = 4.1, *p* = .046, *η*^2^_p_ = .06. The interaction of group and trial type effect was not significant, *F*(1,66) = 2.6, *p* = .11, η^2^_p_ = .04. The P3b was more pronounced after nogo stimuli, *F*(1,66) = 228.4, *p* < .001, *η*^2^_p_ = .78. There was a trend for a group effect, *F*(1,66) = 3.7, *p* = .057, *η*^2^_p_ = .05, and a significant interaction between group and trial type, *F*(1,66) = 7.7, *p* = .007, *η*^2^_p_ = .10. Follow-up analyses indicated that the HBW group showed larger P3b amplitudes than the NBW group on nogo, *t*(66) = 2.5, *p* = .0145 (CI 0.31 2.68), but not on go trials, *t*(66) = 0.65, *p* = .52.

**Figure 2.**
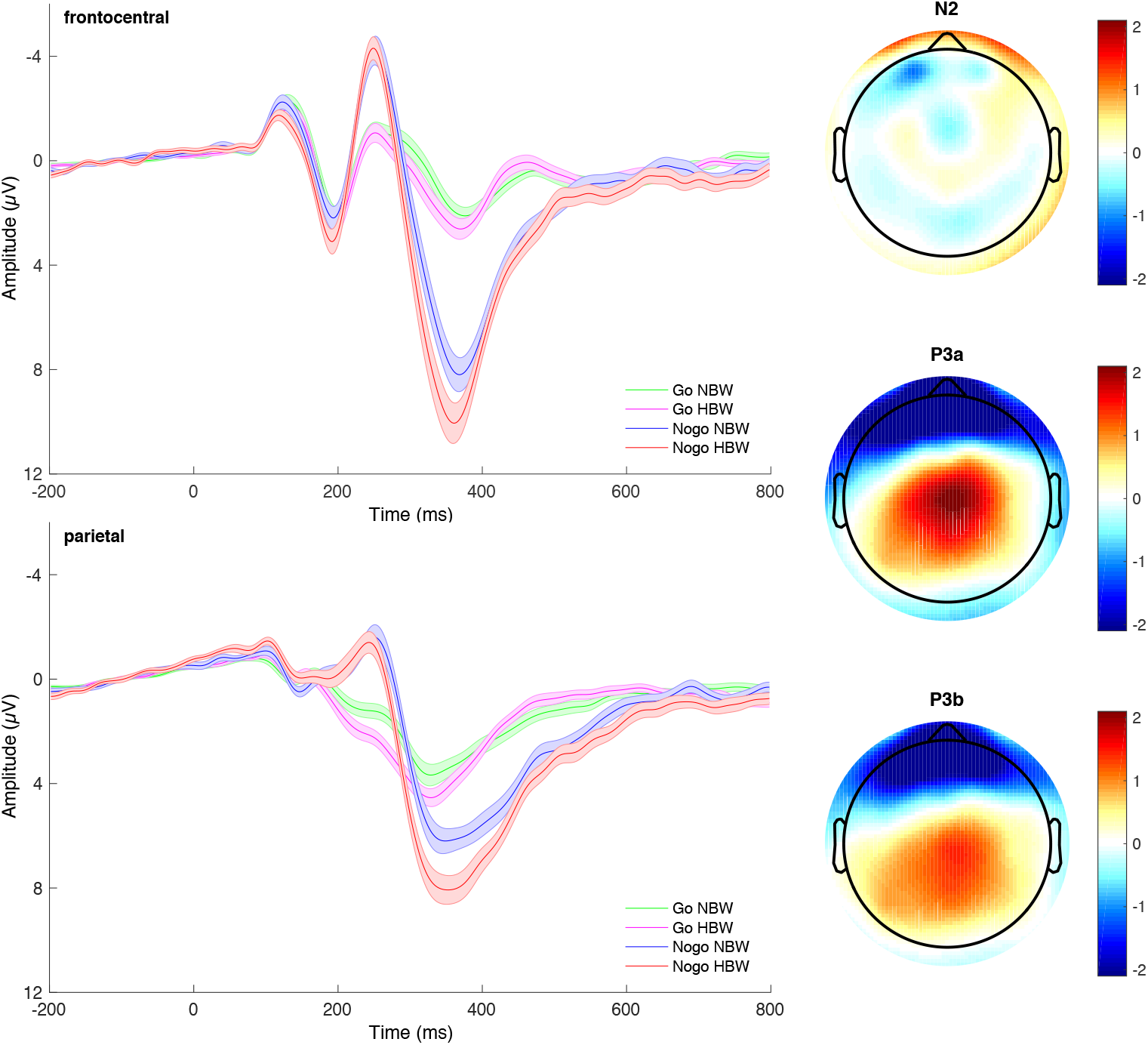
Stimulus-locked ERP waveforms for the high binge watching (HBW) and the no binge watching (NBW) group averaged for go and nogo trials at frontocentral (upper graph) and parietal (lower graph) electrode cluster. Negative values are plotted upward. Solid lines indicate averaged activity and shades represent standard errors. Topographic distribution (top view) are presented in the right column: the difference between HBW and NBW group in N2 (upper graph), P3a (middle graph) and P3b (lower graph) are displayed.

### Neural response: performance monitoring

Group averages and topographies of neural response are displayed in Figure 3. An ANOVA with the factors context, correctness, and group revealed that the negativity (ERN) was significantly larger on error than on correct trials, *F*(1,62) = 241.2, *p* = .001, *η*^2^_p_ = .78. The frontocentral negativity was also more negative in the loss compared to the gain context, *F*(1,62) = 11.6, *p* < .001, *η*^2^_p_ = .16 (*M* = −7.5, *SD* = 3.8 vs. *M* = −6.9, *SD* = 3.9). The main effect of group was not significant, *F*(1,62) = 1.4, *p* < .246, *η*^2^_p_ = .02, but an interaction between group and accuracy was observed, *F*(1,62) = 5.4, *p* = .024, *η*^2^_p_ = .08. Follow-up analyses of the significant interaction were conducted separately for incorrect and correct responses. A significant group main effect was observed following errors, *F*(1,62) = 4.5, *p* = .038, *η*^2^_p_ = .07, but not following correct responses, *F*(1,62) = 0.7, *p* = .396, *η*^2^_p_ = .01. These effects indicate that the amplitude of the ERN was larger for the HBW (*M* = −8.1, *SD* = 3.9) than for the NBW group (*M* = −6.2, *SD* = 3.3). Other significant interactions were not observed.

**Figure 3.**
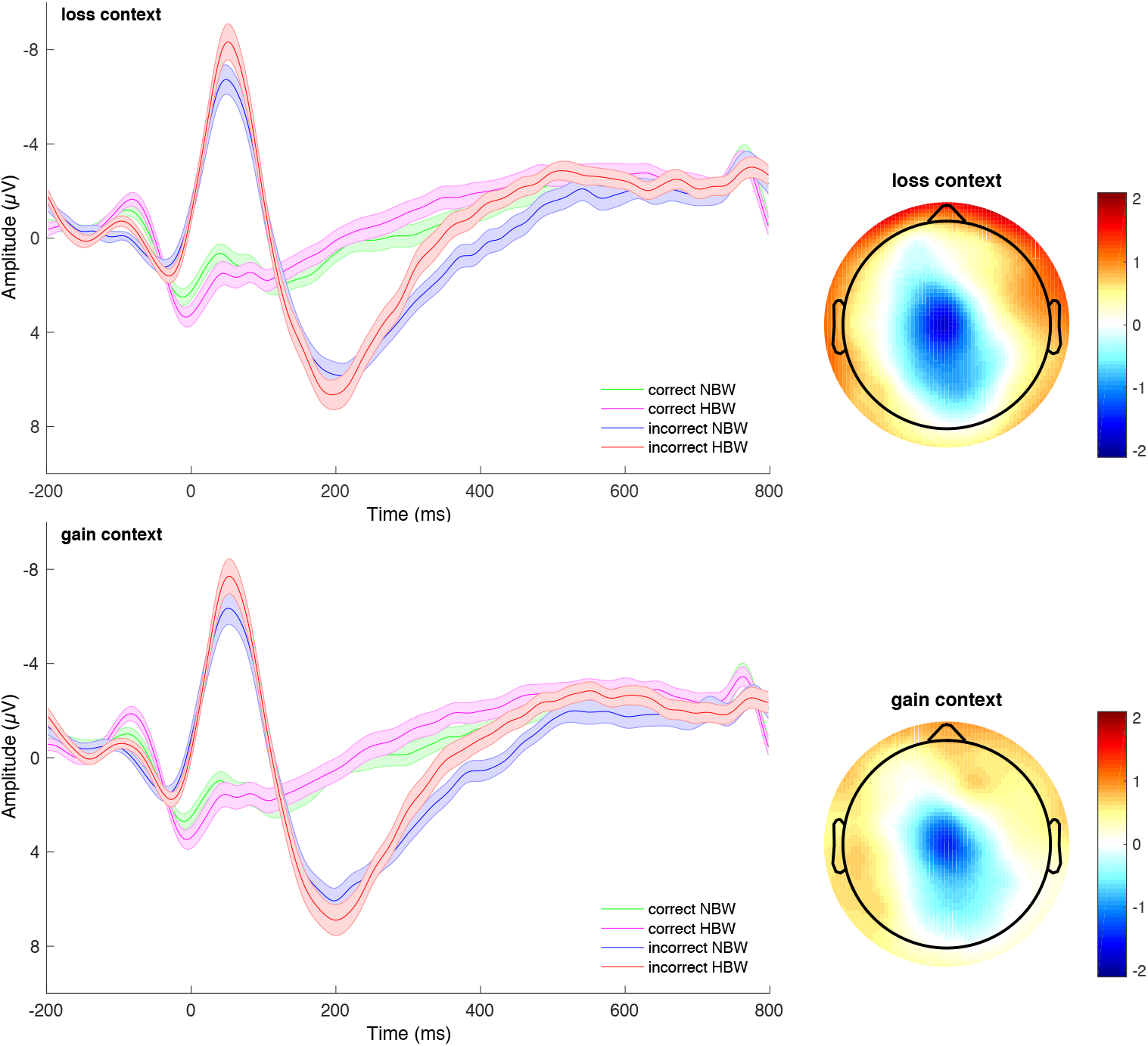
Response-locked ERP waveforms for the high binge watching (HBW) and the no binge watching (NBW) group averaged for correct and incorrect responses at frontocentral electrodes in the loss (top) and gain (bottom) context. Negative values are plotted upwards. Topographic distribution (top view) of the group difference in error-related negativity are presented in the right column: the difference between HBW and NBW group in loss (upper graph) and gain (lower graph) context are displayed.

## Discussion

This is the first study investigating psychophysiological correlates of response inhibition and performance monitoring in individuals with and without binge watching behavior. Both response inhibition and error monitoring were altered in individuals showing binge watching behavior, as indicated by increased amplitudes of P3a, P3b, and ERN.

The P3a or frontal P3 indexes rapid attentional processes and the inhibition of task irrelevant information and the P3b is associated with the inhibition of the response on nogo trials (Polich, 2007). An enhanced nogo P3-complex in the HBW group could represent a correlate of higher neural effort to achieve inhibitory control, to suppress responding, and perform at the same level as the NBW group. If the enhanced inhibitory P3b does reflect such a compensatory mechanism, these findings could suggest reduced inhibitory control in HBW. This may also explain the lack of behavioral differences between the two groups. At the same time, conflict detection, as reflected in the N2, appeared to be unimpaired. This indicates that monitoring processes targeting signals for motor demands (go vs. nogo cues) may function normally in BW, whereas alterations are specific to suppressing motor programs.

This pattern of normal conflict detection and hyperactive response inhibition does not dovetail with inhibition dynamics previously observed in other addictive or excessive behaviors. Whereas internet gaming has also been reported to show unaltered conflict detection (N2; Kim et al., 2017; Li et al., 2019; Littel et al., 2012), response inhibition (P3) was either *reduced* (Li et al., 2019) or *unchanged* (Kim et al., 2017; Littel et al., 2012). Similarly, whereas one study also found increased P3 in the absence of performance differences in excessive internet use and interpreted this finding to reflect cognitive effort when inhibiting successfully (Dong et al., 2010), excessive internet use was repeatedly associated with *impaired* conflict detection (Zhou et al., 2010). The current results do, however, distinctly contrast with findings in excessive smartphone use, for which *increased* conflict detection but *decreased* or *unchanged* response inhibition were reported (Chen et al., 2016; Gao et al., 2019). One implication gained from these inconsistencies is that the spectrum of addictive or excessive behaviors should not be considered homogeneous in terms of processes related to response inhibition. Future research in this young field should move away from comparing smaller groups with high versus low scores in a given construct. Instead, large samples will enable extracting latent classes with distinct inhibitory profiles, which in turn could be linked to expressions in a variety of dimensions of addictive behaviors as well as associated real-life impairments and symptom severity.

Contrasting our hypothesis, the ERN was increased in the HBW group. The ERN is an early signal for performance adaptations after an error occurred (Ullsperger et al., 2014), implying that the HBW group is associated with enhanced performance monitoring. This contrasts with prior reports of reduced ERN amplitudes in addictive behaviors (Franken et al., 2018; Littel et al., 2012; Zhou et al., 2013) and is interesting in light of recent meta-analytic evidence that increased vs. decreased ERN may be a risk-marker for internalizing vs. externalizing disorders, respectively (Pasion & Barbosa, 2019). Provided our ERN result is replicated, this would indicate an internalizing component of BW that seems consistent with the vicious circle account (Panda & Pandey, 2017). Additionally, ERN amplitudes were more pronounced in the loss compared to the gain context which replicates previous studies (Endrass et al., 2010; Potts, 2011; Potts et al., 2006; Riesel et al., 2012). Increased performance monitoring in the face of punishment is in line with findings that motivational factors modulate the performance monitoring system, but it also suggests that cognitive engagement was greater in the face of potential losses relative to gains (Potts, 2011). Although previous research showed that anxiety sensitivity increased and impulsivity decreased (Riesel et al., 2012) the preferential processing of aversive stimuli (Potts et al., 2006), group differences regarding context influences on ERN have not been observed in the current study. However, great caution is advised with this interpretation, particularly given the small effects we observed. In sum, we believe that this surprising result underlines our above-mentioned point to remedy the heterogeneity of ERP results in addictive and excessive behaviors with larger and more meaningful samples.

Limitations of the present study concern sample composition and definition of binge watching. The sample size was too small to allow correlative analyses with enough statistical power. Furthermore, descriptive analyses showed that most participants had high levels of education which is indicative for higher cognitive control and therefore could have led to a bias towards better performance in the behavioral tasks. Furthermore, binge watching and the group classification were defined by the number of binge-watching episodes during a specified period. Psychological symptoms were not considered. Recent studies have shown the usefulness of including psychological symptoms in the definition of binge watching. Therefore, a questionnaire assessing different symptoms related to problematic binge watching should be used (e. g. Series Watching Engagement Scale, Toth-Kiraly et al., 2017). Moreover, it should be considered that binge watching is currently not classified as a mental disorder, even though an addictive potential is debated. Given the lower severity of binge watching, changes on the behavioral or self-report level might not be observable. Finally, behavioral effects for reduced inhibitory performance could not be detected, although the HBW group reported higher impulsivity as measured with BIS-11 score. Also, neither a main effect of context nor an interaction effect of group and motivational context were observed in the MIFLAT.

In conclusion, this study examined neural correlates of cognitive control in binge watching for the first time. Affected individuals showed larger P3a, P3b, and ERN amplitudes compared to individuals without BW behavior. Results indicated poorer response inhibition in BW, similar to findings in behavioral addictions. On the other hand, an altered performance monitoring was observed contradicting the supposed classification. More research is needed to better understand BW behavior and its neuronal correlates.

## Acknowledgements

The authors wish to thank all participants in the study. The authors thank M. Sc. Julia Berghäuser for advice and support regarding the current project. This work was supported by the German Research Foundation (DFG, SFB 940 C6, TRR 265 B1 and EN 906/6-1).

## Authors contribution

TE, CK, KLB, and RO were responsible for the study concept and design. CK and KLB recruited participants and recorded EEG data. CK, KLB, and TE analyzed the data. CK, KLB, RD, and TE were responsible for interpretation. CK and KLB drafted the manuscript and TE, RD, and RO provided critical revisions of the manuscript. All authors critically reviewed the content of the manuscript and approved the final version for publication.

## References

Bari, A., & Robbins, T.W. (2013). Inhibition and impulsivity: behavioral and neural basis of response control. Prog Neurobiol, 108, 44–79.

Chen, J.W., Liang, Y.S., Mai, C.M., Zhong, X.Y., & Qu, C. (2016). General Deficit in Inhibitory Control of Excessive Smartphone Users: Evidence from an Event-Related Potential Study. Frontiers in Psychology, 7.

Delorme, A., & Makeig, S. (2004). EEGLAB: an open source toolbox for analysis of single-trial EEG dynamics including independent component analysis. J Neurosci Methods, 134, 9–21.

Dong, G., Zhou, H., & Zhao, X. (2010). Impulse inhibition in people with Internet addiction disorder: electrophysiological evidence from a Go/NoGo study. Neurosci Lett, 485, 138–142.

Endrass, T., Schuermann, B., Kaufmann, C., Spielberg, R., Kniesche, R., & Kathmann, N. (2010). Performance monitoring and error significance in patients with obsessive-compulsive disorder. Biol Psychol, 84, 257–263.

Enriquez-Geppert, S., Konrad, C., Pantev, C., & Huster, R.J. (2010). Conflict and inhibition differentially affect the N200/P300 complex in a combined go/nogo and stop-signal task. NeuroImage, 51, 877–887.

Euser, A.S., Evans, B.E., Greaves-Lord, K., Huizink, A.C., & Franken, I.H. (2013). Diminished error-related brain activity as a promising endophenotype for substance-use disorders: evidence from high-risk offspring. Addict Biol, 18, 970–984.

Fischer, A.G., Klein, T.A., & Ullsperger, M. (2017). Comparing the error-related negativity across groups: The impact of error-and trial-number differences. Psychophysiology, 54, 998–1009.

Foa, E.B., Huppert, J.D., Leiberg, S., Langner, R., Kichic, R., Hajcak, G., & Salkovskis, P.M. (2002). The Obsessive-Compulsive Inventory: Development and validation of a short version. Psychological Assessment, 14, 485–496.

Folstein, J.R., & Van Petten, C. (2008). Influence of cognitive control and mismatch on the N2 component of the ERP: a review. Psychophysiology, 45, 152–170.

Franken, I.H.A., Nijs, I.M.T., Toes, A., & van der Veen, F.M. (2018). Food addiction is associated with impaired performance monitoring. Biol Psychol, 131, 49–53.

Gao, Q.F., Jia, G., Zhao, J., & Zhang, D.D. (2019). Inhibitory Control in Excessive Social Networking Users: Evidence From an Event-Related Potential-Based Go-Nogo Task. Frontiers in Psychology, 10.

Gehring, W.J., Goss, B., Coles, M.G.H., Meyer, D.E., & Donchin, E. (1993). A neural system for error-detection and compensation. Psychological Science, 4, 385–390.

Goschke, T. (2014). Dysfunctions of decision-making and cognitive control as transdiagnostic mechanisms of mental disorders: advances, gaps, and needs in current research. Int J Methods Psychiatr Res, 23 Suppl 1, 41–57.

Hajcak, G., Moser, J.S., Yeung, N., & Simons, R.F. (2005). On the ERN and the significance of errors. Psychophysiology, 42, 151–160.

Henry, J.D., & Crawford, J.R. (2005). The short-form version of the Depression Anxiety Stress Scales (DASS-21): construct validity and normative data in a large non-clinical sample. Br J Clin Psychol, 44, 227–239.

Huster, R.J., Enriquez-Geppert, S., Lavallee, C.F., Falkenstein, M., & Herrmann, C.S. (2013). Electroencephalography of response inhibition tasks: functional networks and cognitive contributions. Int J Psychophysiol, 87, 217–233.

Kim, M., Lee, T.H., Choi, J.S., Kwak, Y.B., Hwang, W.J., Kim, T., … Kwon, J.S. (2017). Neurophysiological correlates of altered response inhibition in internet gaming disorder and obsessive-compulsive disorder: Perspectives from impulsivity and compulsivity. Scientific Reports, 7.

Koob, G.F., & Volkow, N.D. (2010). Neurocircuitry of addiction. Neuropsychopharmacology, 35, 217–238.

Li, Q., Wang, Y., Yang, Z., Dai, W.N., Zheng, Y., Sun, Y.W., & Liu, X. (2019). Dysfunctional cognitive control and reward processing in adolescents with Internet gaming disorder. Psychophysiology.

Liesefeld, H.R., & Janczyk, M. (2018). Combining speed and accuracy to control for speed-accuracy trade-offs(?). Behavior Research Methods, 51, 40–60.

Littel, M., van den Berg, I., Luijten, M., van Rooij, A.J., Keemink, L., & Franken, I.H. (2012). Error processing and response inhibition in excessive computer game players: an event-related potential study. Addict Biol, 17, 934–947.

Luijten, M., Littel, M., & Franken, I.H. (2011). Deficits in inhibitory control in smokers during a Go/NoGo task: an investigation using event-related brain potentials. PLoS One, 6, e18898.

Luijten, M., Machielsen, M.W., Veltman, D.J., Hester, R., de Haan, L., & Franken, I.H. (2014). Systematic review of ERP and fMRI studies investigating inhibitory control and error processing in people with substance dependence and behavioural addictions. J Psychiatry Neurosci, 39, 149–169.

Olvet, D.M., & Hajcak, G. (2009). The stability of error-related brain activity with increasing trials. Psychophysiology, 46, 957–961.

Orosz, G., Vallerand, R.J., Bőthe, B., Tóth-Király, I., & Paskuj, B. (2016). On the correlates of passion for screen-based behaviors: The case of impulsivity and the problematic and non-problematic Facebook use and TV series watching. Personality and Individual Differences, 101, 167–176.

Panda, S., & Pandey, S.C. (2017). Binge watching and college students: motivations and outcomes. Young Consumers, 18, 425–438.

Pasion, R., & Barbosa, F. (2019). ERN as a transdiagnostic marker of the internalizing-externalizing spectrum: A dissociable meta-analytic effect. Neurosci Biobehav Rev, 103, 133–149.

Patton, J.H., Stanford, M.S., & Barratt, E.S. (1995). Factor structure of the Barratt impulsiveness scale. J Clin Psychol, 51, 768–774.

Polich, J. (2007). Updating P300: an integrative theory of P3a and P3b. Clin Neurophysiol, 118, 2128–2148.

Potts, G.F. (2011). Impact of reward and punishment motivation on behavior monitoring as indexed by the error-related negativity. Int J Psychophysiol, 81, 324–331.

Potts, G.F., George, M.R., Martin, L.E., & Barratt, E.S. (2006). Reduced punishment sensitivity in neural systems of behavior monitoring in impulsive individuals. Neurosci Lett, 397, 130–134.

Riddle, K., Peebles, A., Davis, C., Xu, F., & Schroeder, E. (2018). The addictive potential of television binge watching: Comparing intentional and unintentional binges. Psychology of Popular Media Culture, 7, 589–604.

Riesel, A., Weinberg, A., Endrass, T., Kathmann, N., & Hajcak, G. (2012). Punishment has a lasting impact on error-related brain activity. Psychophysiology, 49, 239–247.

Romer Thomsen, K., Fjorback, L.O., Moller, A., & Lou, H.C. (2014). Applying incentive sensitization models to behavioral addiction. Neurosci Biobehav Rev, 45, 343–349.

Schlienz, N.J., Hawk, L.W., Jr., & Rosch, K.S. (2013). The effects of acute abstinence from smoking and performance-based rewards on performance monitoring. Psychopharmacology (Berl), 229, 701–711.

Sung, Y.H., Kang, E.Y., & Lee, W.-N. (2018). Why Do We Indulge? Exploring Motivations for Binge Watching. Journal of Broadcasting & Electronic Media, 62, 408–426.

Toth-Kiraly, I., Bothe, B., Toth-Faber, E., Haga, G., & Orosz, G. (2017). Connected to TV series: Quantifying series watching engagement. J Behav Addict, 6, 472–489.

Trouleau, W., Ashkan, A., Ding, W., & Eriksson, B. (2016). Just One More. Paper presented at the Proceedings of the 22nd ACM SIGKDD International Conference on Knowledge Discovery and Data Mining - KDD’16.

Ullsperger, M., Fischer, A.G., Nigbur, R., & Endrass, T. (2014). Neural mechanisms and temporal dynamics of performance monitoring. Trends Cogn Sci, 18, 259–267.

Walton-Pattison, E., Dombrowski, S.U., & Presseau, J. (2018). ‘Just one more episode’: Frequency and theoretical correlates of television binge watching. J Health Psychol, 23, 17–24.

Zhou, Z., Li, C., & Zhu, H. (2013). An error-related negativity potential investigation of response monitoring function in individuals with internet addiction disorder. Front Behav Neurosci, 7, 131.

Zhou, Z.H., Yuan, G.Z., Yao, J.J., Li, C., & Cheng, Z.H. (2010). An event-related potential investigation of deficient inhibitory control in individuals with pathological Internet use. Acta Neuropsychiatr, 22, 228–236.

